# Embodied reinforcement learning in the primate cortico-basal ganglia system

**DOI:** 10.64898/2026.07.22.740086

**Authors:** Franco Giarrocco, Bruno B. Averbeck

## Abstract

Learning the value of environmental stimuli from reward experience allows animals to make advantageous choices. Existing biological accounts of reinforcement learning (RL) often assume that this value is represented as a single, motor-system-invariant neural signal. Here we recorded neuronal activity across eight nodes of the macaque cortico-basal ganglia system — from limbic to neocortical regions — while monkeys learned stimulus-reward associations using either saccades or reaches as the required motor response. The geometry of population activity revealed largely distinct value-coding dimensions for saccades and reaches, including in limbic regions typically associated with motor-system-invariant value coding. Consequently, value information was substantially reduced when read out across motor systems. These results provide circuit-wide evidence that challenges existing neural implementations of RL. They link biological accounts of value learning to embodied frameworks in cognitive science and artificial intelligence, in which behavior is grounded in an agent’s physical structure rather than abstracted from it.

## Introduction

Learning which environmental stimuli lead to reward allows animals to make advantageous choices. This process requires animals to build neural representations of stimulus value through reward experience and use these representations to guide future choices. These choices are expressed through motor systems, whose structure and specialization shape how animals interact with their environment. Decades of reinforcement learning (RL) studies in humans and animal models have implicated the cortico-basal ganglia system as a major substrate for these processes ^1–17^, including ventral limbic circuits typically associated with value learning as well as circuits that support oculomotor and skeletomotor functions ^18,19^.

Existing biological accounts of RL often assume that value is represented as a single, motor-system-invariant neural signal. This view is typically formalized in actor-critic models ^5,20–22^. In these models, a critic-like system, associated with ventral striatum and midbrain dopamine neurons, computes a state-value signal, which defines the value of stimuli in the environment, and generates reward prediction errors when outcomes differ from expectation. These errors are thought to provide a common teaching signal that updates both the critic’s value estimate and the actor’s stimulus-response mappings implemented in motor circuits. In this view, value is a unified signal that guides learning across motor systems. This organization is supported by evidence that, under some conditions, midbrain dopamine neurons broadcast a common prediction error throughout the striatum, providing a substrate for shared value updating ^23–28^. Consistent with this, ventral limbic neurons represent value across diverse stimulus types and motor contexts ^29–33^. Stimulus-value coding and stimulus-reward associations also extend to dorsal striatal and neocortical areas ^34–36^, indicating that the motor-system-invariance assumption concerns the broader cortico-basal ganglia system, not ventral limbic circuits alone.

Despite its influence, the assumption of a single, motor-system-invariant value signal has remained largely untested experimentally. Previous studies have investigated the neural substrates of value learning within a single motor system, whether oculomotor or skeletomotor ^29,31,37–40^, precluding a direct test of whether value representations generalize across motor systems. Recent work, however, shows that inactivation of ventral striatum or amygdala impairs stimulus-reward learning differently depending on whether monkeys learn through saccades or reaches ^41^, a result difficult to reconcile with the idea that limbic circuits support value learning through a single motor-system-invariant signal.

Building on principles of embodied cognition ^42–44^, which posit that cognition is grounded in the body’s sensorimotor architecture rather than abstracted from it, we hypothesized that value representations are organized according to the motor system through which learning occurs, an organization we term embodied value. This predicts that value learned through different motor systems should give rise to motor-system-specific value representations rather than being fully abstracted into a single motor-system-invariant signal.

Here, we tested this by recording from 4,843 neurons spanning eight nodes of the cortico-basal ganglia system, from ventral limbic to dorsal cortical areas, while two macaque monkeys learned stimulus-reward associations using either saccades or reaches as the required motor response. The geometry of population activity revealed that value representations were organized along motor-system-specific neural dimensions within this circuit. As a consequence, value information was substantially reduced when read out across motor systems. Within this circuit-wide pattern, separation was especially pronounced in ventral striatum, whereas prefrontal cortex retained relatively more shared geometry across motor systems.

These findings provide circuit-wide evidence that challenges classical neural accounts of RL, revealing value as a representation organized according to the motor systems through which choices are expressed. In doing so, they link biological accounts of value learning to embodied frameworks in cognitive science and artificial intelligence, in which behavior is grounded in an agent’s physical structure rather than abstracted from it ^45–47^.

## Results

### Learning performance and value estimates differ between saccade and reach trials

Two macaque monkeys performed a stimulus-reward learning task, reporting choices with either saccades or arm-reaching movements. The task consisted of randomly intermixed saccade and reach blocks. Each block introduced a new pair of visual stimuli associated with different reward probabilities (80% vs. 20%; Figure 1A). On each trial, monkeys chose one stimulus to receive its associated reward. In saccade trials, monkeys reported their choice with a saccade while keeping the hand on a central target; in reach trials, they reached toward the chosen stimulus while maintaining central fixation.

**Figure 1.**
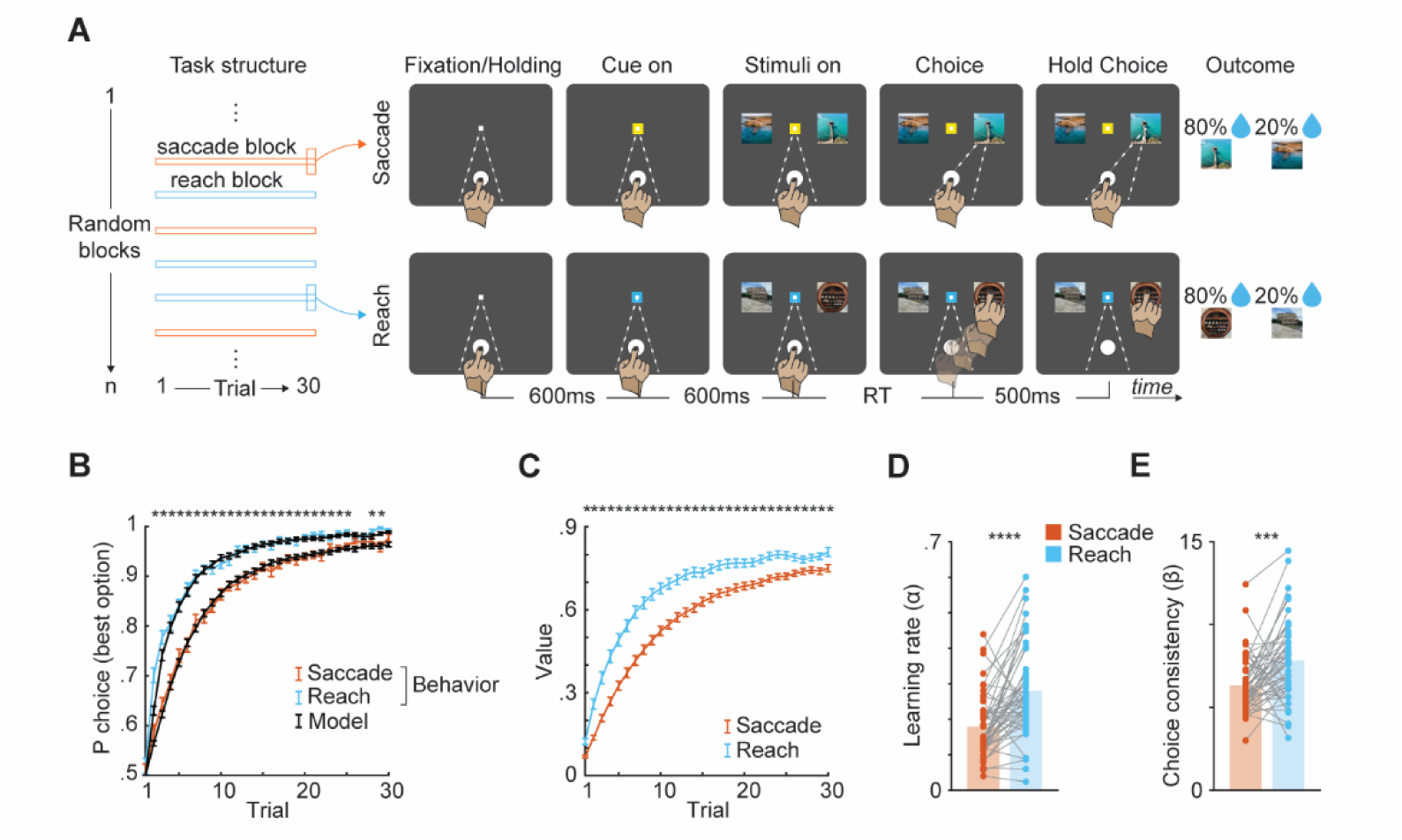
Task design and behavioral results. **A**) Monkeys performed two types of 30-trial blocks randomly intermixed in each session, the saccade and the reach block. In each block, monkeys were presented with a new pair of visual stimuli, each associated with a different probability of reward. Each trial started with the monkeys holding a central target and acquiring a central fixation. Subsequently, a yellow (in saccade trials) or blue (in reach trials) cue appeared around the fixation point, indicating which of the two blocks was in progress. Two stimuli were then randomly presented to either the left or right side of the cue. Monkeys had to choose one of the two stimuli and reward was delivered probabilistically (either 80% or 20% of times) according to the chosen stimulus. Choice was made with a saccade (without detaching the hand) or with an arm-reaching movement (without breaking fixation) toward the chosen stimulus in saccade and reach trials, respectively. **B**) Learning curves. Probability of choosing the most rewarded stimulus across trials for saccade (orange) and reach (blue) blocks. Black lines indicate RL model predictions. **C**) Model estimates of stimulus value across trials for saccade and reach blocks. **D**) Learning rate (mean ± SEM; Saccade = .18 ± .013, Reach = .28 ± .019, paired t-test: t(49) = 5.39, p < .0001) and **E**) choice consistency (mean ± SEM; Saccade = 6.33 ± .24, Reach = 7.83 ± .34, paired t-test: t(49) = 4.04, p < .001) parameters estimated from the RW model, shown for saccade and reach blocks. Individual dots and lines represent individual sessions. Data are pooled across monkeys and reported as mean ± SEM across sessions in which neural activity was recorded (n = 50 sessions). Asterisks indicate significant differences (paired t-test; **** p < 0.0001, *** p < 0.001, * p < 0.05). Supplemental Figure 1 reports results across all behavioral sessions collected, separated by monkey. RT, reaction time.

Learning performance differed across motor systems. Monkeys were more likely to select the higher-value stimulus in reach than in saccade blocks (Figure 1B). To characterize the underlying learning process, we fit a Rescorla-Wagner reinforcement learning model (RW model) to behavior ^48–50^, providing trial-by-trial estimates of the value of each stimulus. The RW model accurately reproduced the monkeys’ choice behavior in both saccade and reach blocks (Figure 1B, black lines). Stimulus-value estimates inferred by the RW model differed between saccade and reach blocks (Figure 1C). This difference was captured by two model parameters: monkeys learned faster in reach blocks, as reflected in a higher learning rate (Figure 1D), and selected the higher-value stimulus more reliably, as reflected in greater choice consistency (Figure 1E). Reach blocks also yielded a higher overall probability of reward (Supplemental Figure 1). These effects were not explained by differences in the number of blocks performed or by measures reflecting differential effort or preference between motor systems, and held across both monkeys over more than one hundred behavioral sessions (Supplemental Figure 1). These behavioral differences between reach and saccade trials are consistent with findings from a previous study investigating explore-exploit behavior, which used a different task structure and allowed unconstrained eye movements during reach choices ^41^. This similarity across studies suggests that the present differences are unlikely to reflect only the specific structure of the present task.

### Single neurons represent value differently in saccade and reach trials

We next asked whether value representations in the cortico-basal ganglia system differed across motor systems. Our dataset comprised 4,843 units recorded from eight nodes of the cortico-basal ganglia system: ventrolateral prefrontal cortex (vlPFC), dorsal premotor cortex (PMd), the caudate (Cd), putamen (Put), medial and lateral ventral striatum (mVS, lVS), the internal globus pallidus (GPi), and the amygdala (Amy) (Figure 2A; Supplemental Figure 2).

**Figure 2.**
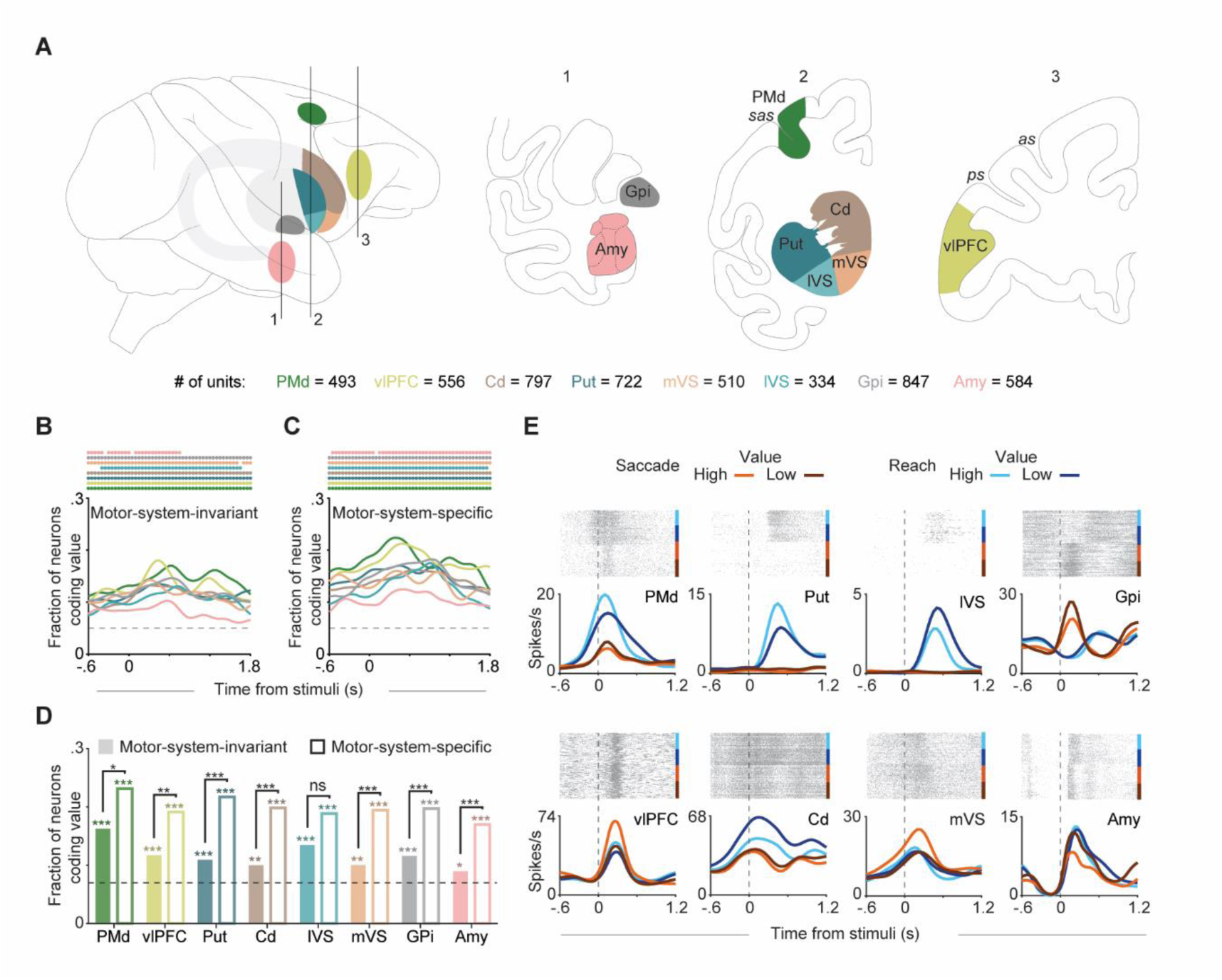
Single-neuron coding of value. **A**) Schematic of recording sites to illustrate sampled areas: dorsal premotor cortex (PMd), ventrolateral prefrontal cortex (vlPFC), caudate (Cd), putamen (Put), medial ventral striatum (mVS), lateral ventral striatum (lVS), internal globus pallidus (GPi), and amygdala (Amy). Numbers indicate total units recorded per region. Supplemental Table 1 reports unit counts by session, region, and monkey. **B**-**C**) Fraction of neurons over time significantly coding value in motor-system-invariant (**B**, main effect of value) and motor-system-specific formats (**C**, value × motor system interaction), shown separately for each region. Colored dots indicate time bins with fraction exceeding chance level (dashed line; binomial test, α = 0.05). **D**) Fraction of neurons with sustained (at least 500 consecutive ms) value coding across regions, shown for motor-system-invariant (filled bars) and motor-system-specific (open bars) coding. Colored asterisks denote fractions significantly above chance (binomial test); black asterisks indicate significant differences between coding types (McNemar test; * p < 0.05; ** p < 0.01; *** p < 0.001; ns, not significant). **E**) Example neurons from each area showing value-related modulation during saccade and reach trials. Raster plots and firing rate (spikes/s) are aligned to stimuli presentation (dashed vertical line). sas, spur of the arcuate sulcus; as, arcuate sulcus; ps, principal sulcus.

We began our analysis by studying single-neuron activity and the extent to which value was represented in a motor-system-invariant format or in a motor-system-specific format. For each neuron, we fit a linear encoding model with stimulus value, motor system identity, choice direction, trial outcome, response time, and their interactions as predictors (see Methods). For each region and time bin, we quantified the fraction of neurons with significant coding of each factor. Neurons with a significant main effect of value showed similar value-related modulation during saccade and reach trials, reflecting motor-system-invariant coding (Figure 2B). Neurons with a significant value × motor system interaction showed value-related modulation that differed between saccade and reach trials, reflecting motor-system-specific coding (Figure 2C).

Under a purely motor-system-invariant value representation, the fraction of neurons with value × motor system interactions should not exceed chance levels. Instead, both motor-system-invariant and motor-system-specific coding exceeded chance levels across regions (binomial test, α = 0.05). Because the task was designed to study how animals learn the value of stimuli over time, each block repeated the same pair of stimuli, making value information continuously available as learning progressed. Accordingly, value coding was present throughout the trial and increased further after stimulus presentation (Figure 2B, C). These results were consistent across both monkeys (Supplemental Figure 3).

To quantify the difference between value coding types, we compared the fraction of neurons with sustained value coding, requiring at least 500 ms of consecutive significance, in either format. Motor-system-specific coding significantly exceeded motor-system-invariant coding across recorded areas (ANOVA, main effect of coding type: F(1,7) = 179.76, p < 0.0001; Figure 2D). This was also true in individual areas, with the only exception in lVS where the difference did not reach significance (McNemar test, p = 0.083). Example neurons from each area illustrate value-related modulation that differed between saccade and reach trials throughout the circuit, including in amygdala and ventral striatum, a pattern that deviates from classical RL predictions (Figure 2E).

### Population geometry reveals embodied value dimensions

Single-neuron analyses revealed the presence of value coding that differed between saccade and reach trials throughout the circuit. However, single-neuron selectivity is inherently ambiguous with respect to the underlying population organization ^51^. Different value modulation between saccade and reach trials could reflect a shared population code with motor-system-dependent gain, consistent with an abstracted organization, or separate population-level representations, consistent with an embodied organization. To distinguish between these possibilities, we turned to a population-geometry approach. Using targeted dimensionality reduction ^50,52^, we identified the population coding dimension tuned to value in each session and area, separately for saccade and reach trials. This dimension is a vector in neural population space capturing the direction along which activity covaries most strongly with stimulus value. To quantify how similar value representations were across motor systems, we measured the angle between the saccade and reach coding dimensions across sessions and areas. Angles were computed in the full 0–180° range, where 0° and 180° indicate fully aligned dimensions (same or opposite sign), and 90° indicates orthogonality, reflecting geometric independence between motor systems. This measure captures both the degree of shared variance and the sign relationship between coding dimensions. The abstracted and embodied hypotheses make distinct predictions (Figure 3A): the abstracted hypothesis predicts aligned dimensions, reflecting a shared value representation, whereas the embodied hypothesis predicts orthogonal dimensions, reflecting distinct value representations.

**Figure 3.**
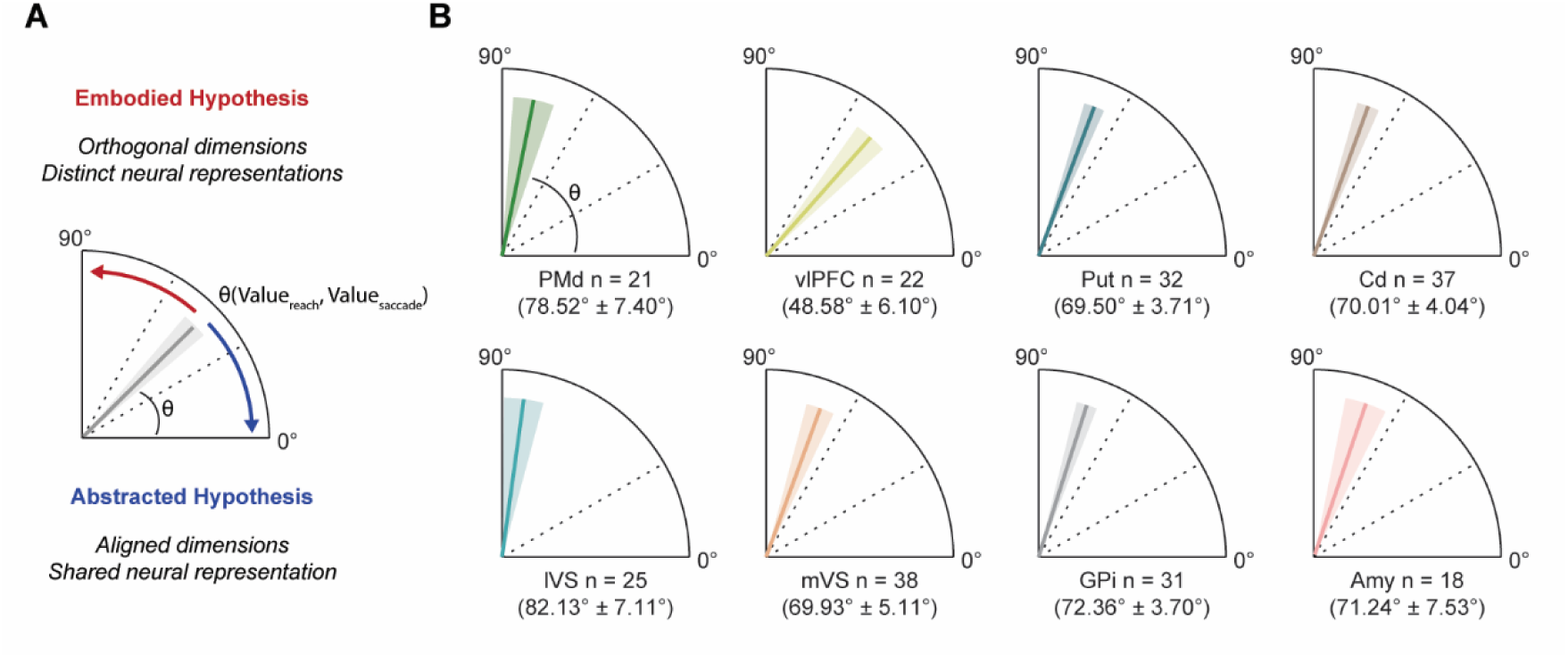
Orthogonality between saccade and reach value dimensions. **A**) Schematic illustration of predictions made by the abstracted and embodied hypothesis. The abstracted hypothesis predicts that value representations are motor-invariant, and therefore saccade and reach coding dimensions (D_reach_ and D_saccade_) should align within neural population space (angle θ approaching 0°), reflecting a shared representation. In contrast, the embodied hypothesis predicts near-orthogonality between saccade and reach coding dimensions (angle θ approaching 90°), reflecting distinct population-level representations. **B**) Angle between saccade and reach coding dimensions in each area. Bonferroni-corrected post-hoc comparisons showed that vlPFC had significantly lower angles than PMd (p = 0.0071) and lVS (p = 0.0011), with marginally lower angles than Put (p = 0.0825), Cd (p = 0.066), mVS (p = 0.0722), and Amy (p = 0.0857). For each area, angles were computed at the single-session level and averaged across sessions (mean ± SEM across sessions; n values indicate the number of sessions per area).

Angles between saccade and reach value coding dimensions were biased toward orthogonality within the cortico-basal ganglia system (mean ± SEM: 70.28° ± 3.50°; Figure 3B), indicating that value representations were organized along largely distinct, motor-system-specific population dimensions rather than along a single motor-system-invariant dimension. Angles varied across areas (ANOVA, main effect of area: F(7,167) = 3.39, p = 0.0021), ranging from 48.58° ± 6.10° in vlPFC to 82.13° ± 7.11° in lVS. Post-hoc comparisons (Bonferroni-corrected) revealed that this effect was driven by vlPFC, which exhibited significantly lower angles than PMd (p = 0.0071) and lVS (p = 0.0011), as well as marginally lower angles than other areas, indicating that vlPFC retained a greater degree of shared geometry between motor systems than other areas. Crucially, neither the medial nor the lateral ventral striatum, which are core components of the critic in actor-critic models, exhibited shared geometry between motor systems.

To assess the robustness of these geometric relationships beyond single-session sampling, we repeated this analysis using pseudo-populations constructed by pooling neurons across sessions within each region. Angles from pseudo-populations confirmed the level of orthogonality observed at the single-session level (Supplemental Figure 4). We further examined whether this geometric separation depended on specific neuronal subpopulations by computing angles separately for neurons with motor-system-specific (significant value × motor system interaction) versus motor-system-invariant (significant main effect of value) coding. Angles remained largely orthogonal when computed from motor-system-specific neurons alone but decreased when restricted to motor-system-invariant neurons, consistent with the expected geometry of each subpopulation (Supplemental Figure 4).

### Embodied geometry constrains cross-system value readout

Because saccade and reach value representations occupy largely orthogonal dimensions, activity patterns along one motor system’s axis carry little information about the other. Therefore, value discriminability should be strongest when neural activity is read out along the matched motor-system dimension (saccade activity along the saccade value dimension, reach activity along the reach value dimension) and should decrease under cross-system readout (saccade activity along the reach dimension, and vice versa). To test this, we constructed a two-dimensional neural space by projecting single-trial activity onto the choice-direction (x-axis) and value (y-axis) coding dimensions, transforming high-dimensional population activity into coordinates along these two task-relevant axes. Incorporating choice direction allowed us to account for the spatial component of the task while isolating value-related modulation. Within this space, neural trajectories tracked the time-resolved mean of projected activity across trials, computed separately for the four combinations of high versus low value and left versus right choice.

When activity was projected onto the matched motor-system value dimension, neural trajectories separated according to both value magnitude and choice direction, with high- and low-value trials diverging along the value axis and left versus right choices segregating along the choice-direction axis (Figure 4A, B, matched). However, value discriminability was reduced in the cross-system neural space (Figure 4A, B, cross). Consistent with an embodied architecture, value discriminability was significantly higher under matched than cross-system readout across regions for both saccade (mean ± SEM Euclidean distance: matched = 1.08 ± 0.13, cross = 0.71 ± 0.11; 35% reduction; ANOVA, main effect of condition: F(1,383) = 62.14, p < 0.0001) and reach trials (matched = 0.93 ± 0.08, cross = 0.65 ± 0.08; 31% reduction; F(1,383) = 62.56, p < 0.0001; Figure 4C, D). These results were consistent in both monkeys (Supplemental Figure 5).

**Figure 4.**
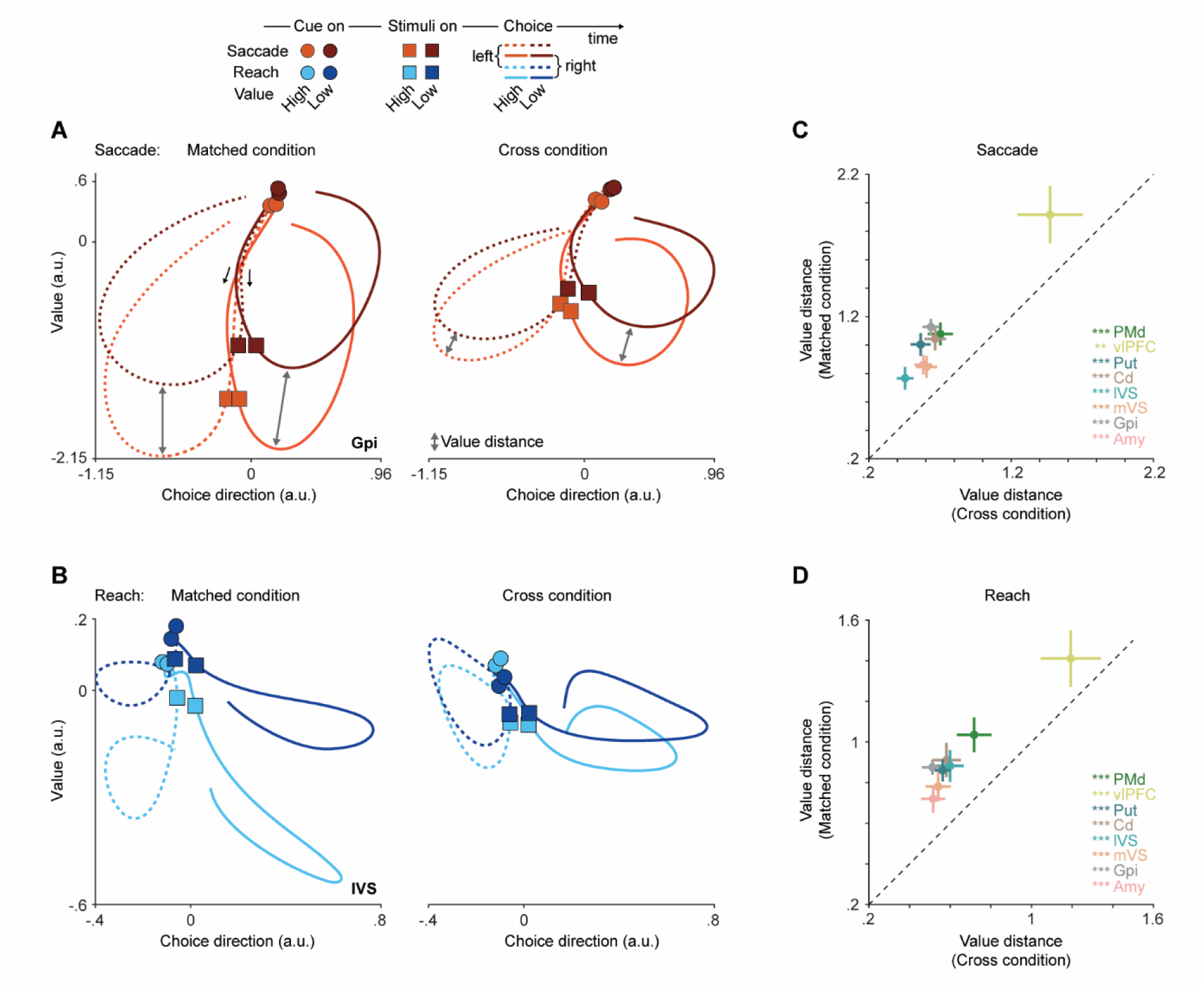
Cross-system value readout. **A**) Two-dimensional neural space for example area (GPi) during saccade trials. Neural trajectories represent low and high value trials, separated by left and right choices, projected onto the choice direction (x-axis) and value (y-axis) coding dimensions. In the matched condition neural space, activity is projected onto the same motor system (saccade) dimensions, whereas in the cross-condition space, activity was projected onto the other motor system (reach) value dimension. Grey arrows depict the distance between high- and low-value trials trajectories (value distance). **B**) same as **A** but for lVS during reach trials. **C-D**) Quantification of value readout during saccade (**C**) and reach (**D**) trials. Each point represents one brain area. Value distance in the matched condition (y-axis) is plotted against value distance in the cross condition (x-axis). The dashed diagonal indicates equality. Points above the diagonal indicate stronger discrimination under matched readout. Error bars indicate ± SEM across sessions. paired t-test: * p < 0.05; ** p < 0.01; *** p < 0.001.

In line with this geometric account, the angle between saccade and reach value dimensions was positively correlated with the reduction in cross-system value discriminability across sessions in all regions, linking the degree of geometric separation to its functional consequence for value readout (Supplemental Figure 6).

Together, these results show that the geometric segregation of value representations constrains value readout across the cortico-basal ganglia system.

### Cross-system decoding reduces value separability

The population geometry analyses revealed that saccade and reach value dimensions were largely orthogonal, predicting reduced cross-system value readout. However, coding angles quantify the relative orientation of value dimensions, not how reliably value can be recovered from population activity because they do not take into account trial-to-trial variability. Two coding dimensions can be aligned geometrically, but if neural activity along them is highly variable across trials, a decoder may not recover value reliably. We therefore trained a linear support vector machine classifier to decode high versus low stimulus value from population activity in each area at the single-session level, providing an independent test of whether value information was organized primarily along motor-system-specific population coding dimensions.

We compared two conditions: within-system decoding, in which a cross-validated classifier was trained and tested on the same motor system, and cross-system decoding, in which the classifier was trained on one motor system and tested on the other. Within-system decoding accuracy was consistently higher than cross-system decoding throughout the trial for both saccade and reach conditions (Supplemental Figure 7). To summarize this difference, we averaged decoding accuracy at the single-session level during the 600 ms following stimulus onset. Within-system decoding accuracy significantly exceeded chance (0.5) across all regions for both saccade and reach trials. Cross-system decoding also exceeded chance, though accuracy was modest, consistent with the degree of orthogonality between saccade and reach value dimensions and the presence of abstracted coding at the single-neuron level. Critically, cross-system decoding accuracy was significantly reduced compared to within-system decoding for both saccade (mean ± SEM accuracy: within = 0.64 ± 0.01, cross = 0.55 ± 0.01; ANOVA, main effect of condition: F(1,383) = 275.73, p < 0.0001; Figure 5A) and reach trials (within = 0.59 ± 0.01, cross = 0.54 ± 0.01; F(1,383) = 168.57, p < 0.0001; Figure 5B). These results were consistent in both monkeys (Supplemental Figure 8). Together with the geometric analyses, these results indicate that value representations across the cortico-basal ganglia system are organized predominantly along motor-system-specific coding dimensions, while retaining a smaller shared component that supports limited cross-system readout.

**Figure 5.**
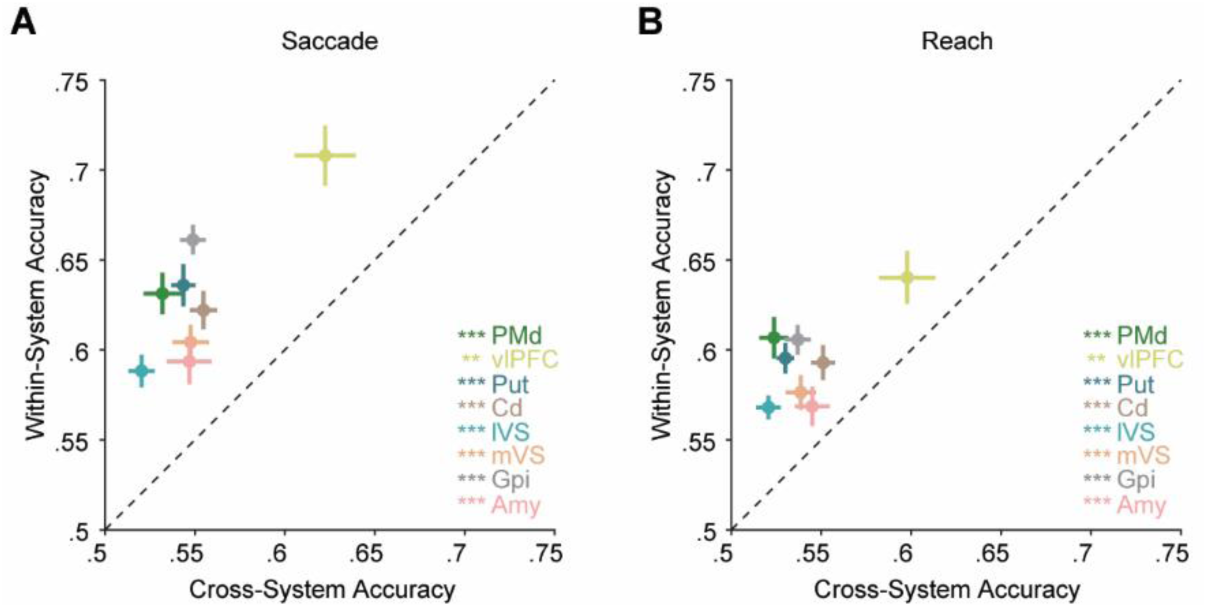
Cross-system decoding of value. **A-B**) Within-system (y-axis) versus cross-system (x-axis) decoding accuracy for high versus low value during saccade (**A**) and reach (**B**) trials. Each point represents the mean ± SEM decoding accuracy for a given area, averaged across sessions during the 600 ms following stimuli presentation. Within-system decoding: classifier trained and tested on the same motor system. Cross-system decoding: classifier trained on one motor system and tested on the other. The dashed diagonal indicates equal decoding accuracy. Areas above the diagonal indicate higher within-system than cross-system accuracy. Colored asterisks denote significant differences between within-system and cross-system accuracy for each area (paired t-test; ** p < 0.01; *** p < 0.001).

### Embodied organization persists under matched behavioral performance

Because learning performance differed between saccade and reach blocks, we next asked whether motor-system-specific value representations could be explained by these behavioral differences. The embodied hypothesis predicts that such representations should persist even in behaviorally matched sessions. We therefore identified a subset of sessions in which learning performance and stimulus-value estimates did not significantly differ between saccade and reach blocks, selected blind to neural activity (Supplemental Figure 9A, B). This yielded eight sessions in which all areas were sampled at least once. We used these sessions to define a binary factor, behavioral similarity (matched vs. divergent behavior), which we introduced into our ANOVAs across the full dataset. Behavioral similarity did not affect the angle between saccade and reach value dimensions (F(1,207) = 0.13, p = 0.72), and this null effect was consistent across areas (behavioral similarity × area: F(7,207) = 1.27, p = 0.27), indicating that the level of orthogonality did not differ in sessions with matched versus divergent behavior. Furthermore, behavioral similarity did not interact with the cross-system effect in either the readout (condition × behavioral similarity: saccade, F(1,429) = 0.16, p = 0.69; reach, F(1,429) = 0.08, p = 0.78) or decoding analyses (saccade, F(1,429) = 0.24, p = 0.62; reach, F(1,429) = 2.90, p = 0.10). Thus, the functional signatures of embodied value were preserved when saccade and reach behavior were matched. Additionally, we applied the population-geometry approach and the cross-system decoding only in sessions with matched behavioral performance, and all results were preserved (Supplemental Figure 9C-E Together, these results indicate that embodied organization does not arise from behavioral differences between motor systems.

## Discussion

Our findings reveal that when value is learned through distinct motor systems, it is represented in largely separate neural spaces across the cortico-basal ganglia circuit, rather than in the motor-system-invariant format assumed by classical biological accounts of RL. Although some single cells showed motor-system-invariant representations, most neurons showed motor-system-specific value coding, and population activity was organized into segregated value coding dimensions for saccades and reaches. As a result, value information was substantially reduced when read out across motor systems.

The degree of geometric separation of value varied across areas, providing insight into the circuit-wide architecture of value learning. vlPFC exhibited the greatest degree of shared geometry between motor systems, consistent with its role in representing abstract task variables ^53–56^. vlPFC receives dominant inputs from temporal cortical visual areas, where stimulus identity is represented in an effector-independent format, but also connects to premotor areas, which in our data showed predominantly motor-system-specific value coding ^57–59^. vlPFC may therefore serve as an interface where motor-system-invariant stimulus representations intersect with motor-system-specific value codes, maintaining a more abstracted value representation while the broader circuit shows greater motor-system-specific organization.

If vlPFC represents one end of this organization, retaining more shared geometry, the ventral striatum represents the other. The lVS, in particular, approached full orthogonality. Crucially, this is the observation most in tension with classical actor-critic accounts because a region typically associated with a critic-like role showed the strongest geometric segregation of value.

Classical RL formalisms distinguish two targets of value learning: state value, the value of the current configuration of the environment, and action value, the value of a specific action in a given state ^3,60^. These values can coincide or diverge depending on task structure. In our task, the state on each trial is defined by the spatial arrangement of the presented stimuli and is independent of the action taken on the previous trial. There are therefore no action-state transitions, and the state of the environment is fully specified by the identity of the presented stimuli. A standard centralized actor-critic account would therefore predict a single, motor-system-invariant stimulus-value signal in the ventral striatum, a prediction challenged by the geometric segregation observed here.

Causal evidence from lesion studies further constrains actor-critic frameworks. Ventral striatal lesions selectively disrupt stimulus-based but not action-based learning ^49^, a dissociation that appears consistent with a critic-like role for ventral striatum. This account, however, leaves two key observations unexplained. First, it does not predict that the identity of the motor system used to express behavior should influence learning. Second, if ventral striatum provides a common critic-derived teaching signal for updating action policies, then disrupting this structure should also impair action-based learning, yet action-based learning was preserved. Furthermore, lesions of the ventral striatum and amygdala impair stimulus-reward learning in some tasks ^48,61^ but not others ^62,63^, suggesting that the contribution of these structures to learning is conditional on task demands. Importantly, we previously showed that when the task requires learning through different motor systems, these lesions produce different effects ^41^, providing causal evidence that the critic’s contribution to value learning is not motor-system-invariant.

Beyond the value representations themselves, the assumption that dopamine neurons broadcast a uniform reward prediction error across striatal targets and provide a common substrate for value updating ^23,24^ has also come under reconsideration. Recent work has shown that dopamine signals are more heterogeneous than initially assumed ^64–67^, with neuronal populations encoding functionally dissociable signals ^68–71^, and regionally heterogeneous release patterns and connectivity that deviate from classical RL predictions ^67,72–75^. Together with our results, this convergence challenges the assumption of a single, motor-system-invariant value-learning system.

The saccade and reach systems also showed different learning performance. One possible explanation is that the two systems play complementary ecological roles. Primates rely heavily on vision to gather information about their environment, an adaptation reflected in the expansion of visual cortical areas across primate lineages ^76–78^. In many contexts, the oculomotor system provides a rapid means of orienting to and sampling behaviorally relevant objects ^79^, whereas the skeletomotor system enables direct physical interaction with those objects to achieve goals. The difference in learning performance observed here may therefore reflect the distinct ecological functions and sensorimotor demands of the two systems. Consistent with this interpretation, we previously found that saccade behavior favored exploration whereas reach behavior favored exploitation and yielded better learning performance in a distinct explore–exploit task ^41^. Because that prior task differed from the present design, including by allowing unrestricted eye movements during reaching, the convergence across studies suggests that the behavioral asymmetry observed here is not simply an artifact of the current task structure.

This behavioral divergence does not, however, account for the geometric separation we observed. We found that value representations remained geometrically segregated when behavioral performance was matched between saccade and reach systems, supporting the idea that embodied organization reflects a property of the circuit rather than a byproduct of performance differences.

If value is organized within each motor system rather than shared between them, what functional advantage might this confer? Animals often interact with their environment through multiple motor systems simultaneously. A primate, for example, may monitor predators with gaze while using its limbs to climb. In social contexts, gaze can also signal attention or intention to others, while the limbs are engaged in separate actions. Such scenarios impose simultaneous and often different demands on distinct motor systems ^80^. A single abstracted value signal shared across systems would have to support choices across action spaces with very different ecological and biomechanical demands. An embodied architecture, by contrast, would allow each motor system to maintain value estimates calibrated to its own action space. This view aligns with the affordance competition hypothesis ^81^, in which potential actions compete for selection based on their sensorimotor specifications, a competition that motor-system-specific value signals could directly support. The largely orthogonal geometry we observed would reduce interference between motor systems, with the value representation of one system insulated from the value estimates of another. At the same time, the angles between value coding dimensions approached but did not reach complete orthogonality, and cross-system decoding remained modestly above chance. This residual sharing could provide a channel for coordination when behavior requires it, while the dominant orthogonal structure keeps each system’s value representation distinct.

Motor system evolution has shaped not only the repertoire of actions available but also the architecture of the cognitive systems that use them ^82^. This is central to embodied cognition frameworks, in which cognitive processes are grounded in the sensorimotor structure of the body rather than fully abstracted from it ^42–44,83^. Embodied decision frameworks apply this principle to decision-making, proposing that action is not merely the output of a completed decision but an integral part of the decision process itself ^84–86^. Consistent with this view, work on pragmatic coding in cortico-striatal circuitry shows that representations of stimuli and goals are formatted according to their behavioral relevance and the actions they predict ^87–89^. Our findings place value representations within this embodied framework: when monkeys learned stimulus-reward associations through saccades or reaches, value was organized along largely motor-system-specific dimensions within the same recorded neural populations.

Beyond biological accounts of value learning, our findings parallel a shift in artificial intelligence toward modular and embodied architectures. Early work in robotics argued that adaptive behavior emerges from direct sensorimotor coupling with the environment rather than from abstracted, centralized representations ^90^. Building on this, the field of AI and robotics has moved toward biologically inspired architectures in which behavior is grounded in the physical structure of the agent ^45,46^. These approaches have been applied effectively in robotics, where agents coordinate multiple actuators with different physical properties and action spaces, and where modular architectures have been favored over centralized control schemes ^45,91^. Our findings provide biological evidence for a similar organizational principle.

Our findings invite comparison with the common currency hypothesis, which posits that values are represented in a shared reference frame to enable comparison across options during economic choice ^92^. The two frameworks operate at distinct stages of value-guided behavior. Common currency concerns how heterogeneous subjective utilities are mapped onto a shared scale at the moment of choice, when options must be compared, value is subjective, and there is therefore no a priori better choice ^93^. Embodied value, in contrast, concerns how value is acquired during reinforcement learning, where reward contingencies are learned through experience and, like in the present task, there is an objectively better choice. The two accounts are thus complementary rather than competing: embodied value describes how values are acquired within motor-system-specific representations, whereas common currency describes how values can be compared across options at the moment of choice.

Finally, our results may also be relevant to psychiatric conditions in which reinforcement learning is disrupted ^94,95^. In obsessive-compulsive disorder and substance use disorders, maladaptive behaviors are often expressed through specific behavioral routines, such as compulsive rituals or drug-seeking actions ^96,97^. If value representations are organized along motor-system-specific dimensions, disruptions to these representations may manifest within particular action contexts rather than uniformly across behavior. This perspective is consistent with the efficacy of behavioral therapies that directly target the specific behavioral routines through which symptoms are expressed ^98,99^. Whether such symptoms are associated with altered geometry of value representations across motor systems remains an important question for future work.

In conclusion, the present findings extend the view of value during reinforcement learning from a single internal quantity computed centrally and then read out by motor systems to a representation that can be organized according to the motor systems through which choices are expressed, at least when learning proceeds within distinct effector systems. In doing so, they bring the neural substrate of reinforcement learning into closer alignment with embodied accounts of cognition across biology and artificial intelligence.

### Limitations

Our task isolates learning within each motor system in alternating blocks, and therefore does not test whether the geometric separation we observe is maintained, strengthened, or reorganized when saccade and reach learning proceed simultaneously. We investigated value learning in the oculomotor and skeletomotor systems; whether these findings extend to other effectors, such as bimanual coordination or vocalization, or to species with different sensorimotor repertoires, remains an important question for future work. Evidence from humans supports this possibility: in a simultaneous bimanual reinforcement-learning task, behavior was better explained by a model that decomposed values into effector-specific components than by a model treating bimanual actions as a unitary choice ^100^, suggesting that effector-specific value learning can operate beyond the saccade-reach distinction.

## Methods

### Subjects

Two 9-year-old male rhesus monkeys (Macaca mulatta) were used in this study. Animals were maintained under controlled water access and earned fluids through task performance on testing days. All procedures conformed to the *Guide for the Care and Use of Laboratory Animals* and were approved by the NIMH Animal Care and Use Committee.

### Task design and stimuli

Monkeys performed the behavioral task while sitting in a primate chair inside a dark, acoustically isolated room with their heads restrained. Visual stimuli were presented on a 19-inch LCD touchscreen monitor (1915L Elotouch) situated 40 cm from the monkeys’ eyes. Stimulus presentation and behavioral monitoring were controlled by MonkeyLogic ^101^. Eye movements were tracked using the ViewPoint EyeTracker system (Arrington Research) at a sampling rate of 1 kHz. On rewarded trials, a fixed volume of apple juice was delivered via a pressurized system controlled by a solenoid valve ^102^.

The task consisted of two types of 30-trial blocks that were randomly intermixed within each session, the saccade and the reach blocks. Each trial began once the monkey touched a central touch target and directed gaze toward a fixation point located 8° of visual angle above the central touch target; both hand contact and fixation were then maintained for 600 ms. Subsequently, a square cue surrounding the fixation point appeared on every trial indicating the current block type. A yellow and blue square signaled saccade and reach blocks, respectively. After 600 ms, two images (referred to as visual stimuli, or stimuli) were presented randomly to the left (−8°) and right (+8°) of the cue. The monkey was required to choose one of the two stimuli, each associated with a different probability of reward. In each block, a novel pair of stimuli was introduced; one stimulus was assigned a high reward probability (p = 0.8) and the other a low reward probability (p = 0.2), with assignment randomized across blocks. In saccade trials, the monkey indicated the choice by making a saccade toward the selected stimulus while holding the central touch target. In reach trials, the monkey indicated its choice with a reaching movement toward the selected stimulus while maintaining central fixation. The monkey was free to make the choice at any time after the stimuli presentation. The monkey was required to hold the choice for 500ms, without detaching the hand from the central touch target in saccade trials, and without breaking fixation in reach trials. Following this 500 ms period, the monkey received the reward according to the probability associated with the choice. Each trial ended with a 500 ms period during which the monkey was free to view both images, followed by a 1500ms black screen intertrial time interval (ITI). During reach trials, breaking fixation at any time before outcome resulted in an error. Similarly, during saccade trials, detaching the hand from the central touch target at any time before outcome resulted in an error. Not holding the choice for the required 500ms also resulted in an error. Errors immediately terminated the trial and initiated the ITI. The location of stimuli was randomized in every trial, including trials following an error.

Each session included four additional blocks (two saccade and two reach blocks) in which a set of four images with known reward probabilities were presented. These trials were excluded from all analyses.

Images were normalized for luminance and spatial frequency using the SHINE toolbox for MATLAB, as described previously ^61^.

### Surgical procedures

Each monkey was surgically implanted with a titanium headpost and a custom 25 × 30 mm recording chamber to permit vertical grid access to the targeted recording sites. Burr holes were drilled to provide access to the brain. Guide tubes were inserted through the recording grid and advanced through the burr holes toward the targeted areas. For subcortical recordings, guide tubes were positioned approximately 5 mm above the target structure. For cortical recordings, guide tubes were advanced to approximately 1.5 mm below the dura. After placement, guide tubes were secured to the grid. Guide tubes were replaced every 5-7 recording days by inserting new tubes at adjacent grid locations. Chamber placement and guide tube trajectories were planned and verified using T1- and T2-weighted magnetic resonance imaging (Supplemental Figure 2). All surgical procedures and MRI scans were performed under sterile conditions and general anesthesia.

### Neurophysiological recordings

Neurophysiological recordings began after postoperative recovery. On each recording day, linear multielectrode probes (V-probe; Plexon Inc., Dallas, TX) were lowered through the guide tubes until reaching the target area. 32-channel electrodes with 150 μm inter-contact spacing probes were used in the PMd, vlPFC, and GPi. 64-channel electrodes with 150 μm inter-contact spacing probes were used in the lateral anterior striatum (simultaneously targeting the Put and lVS), in the medial anterior striatum (simultaneously targeting the Cd and mVS), and in the Amy. On each recording day, we acquired signal from up to 5 simultaneous probes. The probes were advanced to target depth using an 8-channel micromanipulator (NAN Instruments, Nazareth, Israel) attached to the recording chamber. Electrode depth was estimated relative to the guide tube tip position, verified with MRI. vlPFC recordings spanned areas 46v, 8a, 45, and the most dorsal portion of area 12l, between +30 mm and +35 mm along the antero-posterior axis in stereotaxic coordinates. PMd recordings spanned the ventro-rostral portion of area F2 around the spur of the arcuate sulcus and the dorsal portion of the arcuate sulcus. Anterior striatal recordings extended from +22 mm to +28 mm along the antero-posterior axis. Within the striatum, neurons ventral to the internal capsule were classified as mVS or lVS based on their estimated recording locations, determined from MRI-based localization of the guide tube tip and the known inter-contact spacing along the probe. The two populations were not pooled, as their response patterns differed. Amygdala recordings targeted the basolateral nucleus.

Electrophysiological signals were acquired with a 512-channel Grapevine System (Ripple, Salt Lake City, UT). The spike detection threshold was set at a 4.0 × root mean square (RMS) of the baseline signal for each electrode. Behavioral event markers from MonkeyLogic and eye-tracking signals from Viewpoint were sent to the Ripple acquisition system. Extracellular signals were digitized at 30 kHz for single-unit recording. Spike sorting was performed offline using Offline Sorter (Plexon Inc.). Comprehensive methodological details, including components of the recording chamber, guide tube insertion procedures and assembly, have been described in our previous study ^50^.

### Behavioral data analysis

Sessions comprised multiple 30-trial blocks in which monkeys chose between two stimuli associated with either 80% or 20% reward probability. We measured choice behavior across blocks by aligning trials by their position within each block and calculating, for each trial (1-30), the proportion of blocks in which the higher-reward-probability stimulus was selected.

To characterize the learning process, we fit RL models to monkeys’ behavior. Specifically, we fit a variant of the Rescorla-Wagner model to estimate the value of the chosen stimulus as well as the learning rate and inverse temperature, as in previous studies ^48–50^. Stimulus value updates at each trial, t, were given by:

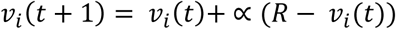

where *v_i_* is the value estimate for stimulus *i*, *R* is the reward feedback for the current choice (1= rewarded, 0 = non rewarded), and α is the learning rate parameter. These value estimates were then passed through a logistic function to generate the choice probability estimate for each stimulus on each trial:

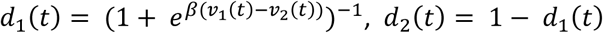

Where β is the inverse temperature parameter or choice consistency. and *d_1_* and *d_2_* are the two choice probability estimates for the stimulus 1 and 2, respectively. We then maximized the likelihood of the monkeys’ choices, *D*, given the parameters, using the following function:

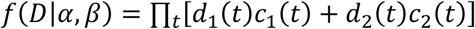

where *c*_1_(*t*) had value of 1 if option 1 was chosen on trial *t*, and *c*_2_(*t*) had value of 1 if option 2 was chosen on trial *t*. Otherwise, they had a value of 0.

Only completed trials were included in the analysis. Trials in which the monkey broke fixation, detached the hand, failed to make a choice, or attempted to saccade or reach toward more than one target were excluded. Trials in which the monkey did not engage in the task were rare events (≤ 0.02% of trials in every session) and were also excluded from the analysis.

### Neuronal encoding of value

To quantify value coding and its relationship to motor systems, we fit a linear model to each neuron’s activity using task and behavioral variables as well as their interactions as predictors of the firing rate. These variables included the stimulus value estimate from the model, motor system identity (whether the trial was a saccade or reach trial), choice direction (whether the monkey selected the left or right stimulus), trial outcome (rewarded or not rewarded), and response time (the time between the stimuli presentation and the monkey’s response). Including all relevant variables in a single model allowed us to isolate value-related encoding from the contribution of other task-related signals. We computed the firing rate of each neuron in 250-ms windows, advanced every 50 ms, from −1.2 s to 1.8 s relative to stimuli presentation, at the single-trial level. We z-scored the activity of each neuron throughout the entire session. The model was fit to all trials jointly, including both saccade and reach trials.

This approach decomposed each neuron’s activity into components shared between motor systems (main effects) and components that differed between motor systems (interaction terms). We defined abstracted value neurons as those showing a significant main effect of value, reflecting value encoding invariant to the motor system. We defined embodied value neurons as those showing a significant value × motor system interaction, reflecting value encoding that differed as a function of the motor system. Significance for main effects and interactions was assessed independently at each time bin based on the p-value of the corresponding regression coefficient (p < 0.05). Fractions of coding neurons were considered significantly higher than expected by chance using a binomial test (α = 0.05). To identify neurons with sustained coding, we required significance in at least two consecutive non-overlapping 250-ms bins (≥ 500 ms) within the analysis window. Full results for all main effects and interactions are reported in Supplemental Figure 3. We then tested whether the observed fractions were larger than expected by chance through a binomial test. The chance level for the binomial test was computed analytically as the probability that a neuron would show at least two consecutive significant non-overlapping bins by chance given the number of bins tested within the analysis window and a per-bin false-positive rate of α = 0.05. To compare the prevalence of abstracted and embodied coding within each area, we used McNemar’s test on paired classifications (abstracted vs. embodied) for each neuron. Across areas, we tested whether embodied coding exceeded abstracted coding using a two-way ANOVA with coding type (embodied vs abstract) and area as factors.

### Targeted dimensionality reduction and population geometry

To characterize the population-level organization of value representations across motor systems, we fit a linear model to each neuron’s activity separately for saccade and reach trials at the single-session level. As in the joint model, we included all task and behavioral variables (stimulus value, choice direction, outcome, and response time) as predictors, allowing us to isolate value-related encoding from other task-related signals. From this model, we applied targeted dimensionality reduction ^50,52^ to derive a population coding dimension for each task variable within each session and region.

For each area and variable, the regression yielded a matrix of coefficients (neurons × time bins) reflecting the strength of encoding over time. We identified the time bin at which the vector of coefficients across neurons had the largest Euclidean norm. The corresponding vector (one coefficient per neuron) was taken as the population coding dimension for that variable, defining an axis in neural population space along which activity varies most strongly with the respective task variable. We derived distinct coding dimensions for stimulus value and choice direction, computed independently for saccade and reach trials.

To quantify the similarity between saccade and reach coding dimensions for each variable, we computed the angle (θ) between their respective population vectors at the single-session level. Saccade and reach coding dimensions were first normalized to unit length, and the angle was obtained from their dot product using the inverse cosine function, yielding values in the full 0°–180° range, where 0° and 180° indicate fully aligned dimensions (same or opposite sign, respectively) and 90° indicates orthogonality.

To test whether this geometric separation had functional consequences for value readout, we constructed a two-dimensional neural space by projecting single-trial activity onto the choice-direction and value coding dimensions. Within this space, neural trajectories were computed as the time-resolved mean of projected activity for high- and low-value trials, separated by left and right choices. High and low value trials were defined based on the 33rd and 66th percentiles of the value distribution.

For each motor system, we constructed two neural spaces: a matched space, in which single-trial activity was projected onto the choice and value dimensions of the same motor system (e.g., saccade trials onto saccade dimensions), and a cross space, in which activity was projected onto the value dimension of the other motor system while retaining the choice dimension of the same motor system (e.g., saccade trials onto the saccade choice dimension and the reach value dimension). Value discriminability was defined as the mean Euclidean distance between high- and low-value trajectories, computed within each choice direction. This analysis was performed separately for each area and motor system at the single-session level. To compare matched-system and cross-system value readout, we used an ANOVA with condition (within, cross) and area as factors, and monkey and session as blocking factors, performed separately for saccade and reach trials. Post-hoc comparisons were performed using paired t-tests with Bonferroni correction. To assess the relationship between geometric separation and value readout, we computed, separately for each area, the Pearson correlation between the angle separating saccade and reach value dimensions and the reduction in cross-system value discriminability (defined as the difference between matched and cross-system Euclidean distances) across sessions.

### Cross-system decoding of value

To provide an independent measure of the value information accessible from population activity, we trained a support vector machine (SVM) classifier to decode high versus low stimulus value from z-scored population activity. As for the cross-system readout analysis, high and low value trials were defined based on the 33rd and 66th percentiles of the value distribution. The classifier was applied to activity in 250-ms windows advanced every 50 ms, at the single-session level for each area.

We compared two conditions. In within-system decoding, the classifier was trained and tested on trials from the same motor system using 10-fold cross-validation. In cross-system decoding, the classifier was trained on all trials from one motor system and tested on all trials from the other. Decoding accuracy was computed at each time bin. For summary analyses, we averaged decoding accuracy of each session during the 600 ms following stimuli presentation.

To compare within-system and cross-system decoding accuracy, we used an ANOVA with condition (within, cross) and area as factors, and monkey and session as blocking factors, performed separately for saccade and reach trials. Post-hoc comparisons were performed using paired t-tests with Bonferroni correction.

## Acknowledgments

We thank Andrew Mitz, Marta Andujar, and Diana Burk, for valuable discussions and comments on the manuscript. We thank Ramon Bartolo, Andrew Mitz, and Diana Burk for technical and surgical help. Anatomical MRI scanning was carried out in the Neurophysiology Imaging Facility Core (NIMH, NINDS, NEI). This research was supported [in part] by the Intramural Research Program of the National Institutes of Health (NIH) (ZIAMH002928 to B.B.A.), a NARSAD Young Investigator Grant from the Brain & Behavior Research Foundation (32825) to F.G., and by the Center on Compulsive Behaviors, NIH via the NIH Shared Resource Subcommittee to F.G.. The contributions of the NIH author(s) are considered Works of the United States Government. The findings and conclusions presented in this paper are those of the author(s) and do not necessarily reflect the views of the NIH or the U.S. Department of Health and Human Services.

## Author contributions

F. G. and B.B.A. designed the study; F.G. performed the experiment; F.G. and B.B.A analyzed the data; F.G. wrote the manuscript; F.G. and B.B.A. revised the manuscript. F.G. and B.B.A. acquired funding.

## Data availability

Custom code used to generate the results reported in this study is publicly available at https://github.com/GiarroccoFranco/Cortico-BasalGanglia.

The data supporting the findings of this study will be made publicly available in a suitable repository upon publication. Prior to publication, the data are available from the corresponding authors upon request.

## Materials & Correspondence

Correspondence and material requests should be addressed to F.G.

