## Supplemental Figure 1-9 and Supplemental Table 1 for "Embodied reinforcement learning in the primate cortico-basal ganglia system"

**Supplemental Figure 1-9; Supplemental Table 1**

***Corresponding Author**
Franco Giarrocco, Ph.D.
Laboratory of Neuropsychology, NIMH/NIH
Building 49 Room 1B80
49 Convent Drive MSC 4415
Bethesda, MD 20892-4415


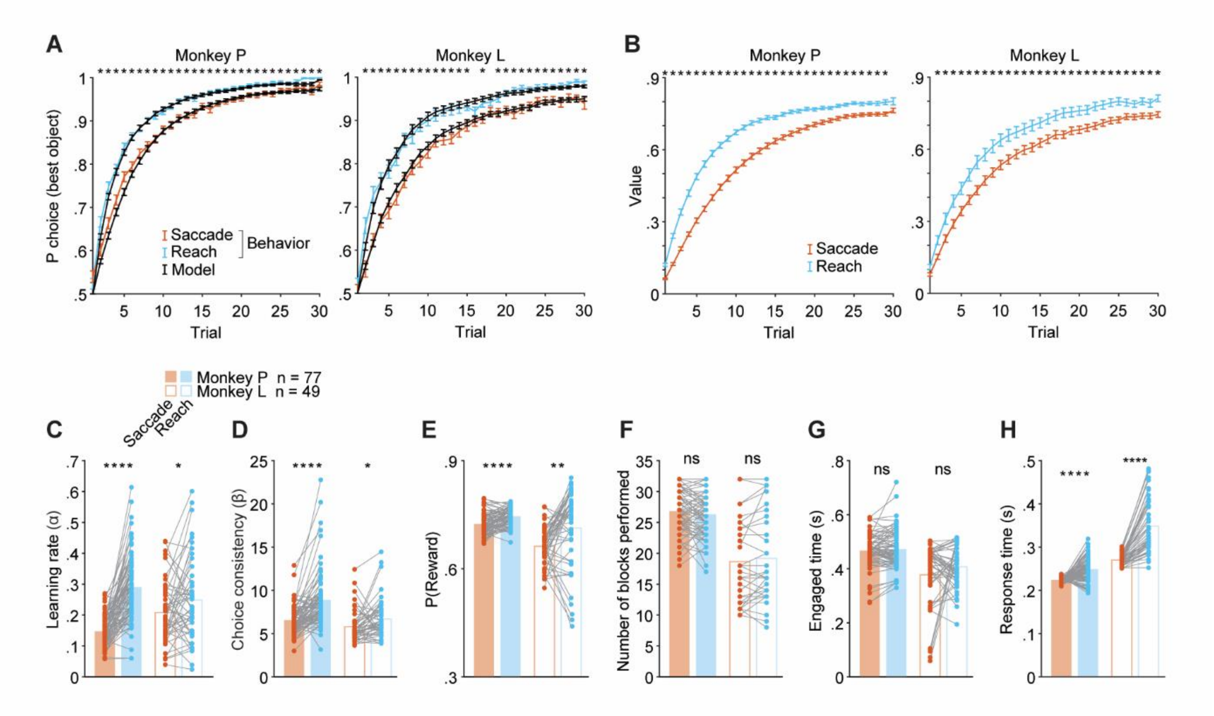


**Supplemental Figure 1. Behavioral and RL model results separated by monkey**. **A**) Learning curves. Probability of choosing the higher-value stimulus across trials for saccade (orange) and reach (blue) blocks. Black lines indicate RL model's choice predictions. Data are reported as mean ± SEM across sessions (Monkey P: n = 77; Monkey L: n = 49). Asterisks indicate significant differences between saccade and reach blocks (paired t-test, $p < 0.05$). **B**) Model estimates of stimulus values across trials for saccade and reach blocks. **C–H**) Model and behavioral parameters shown separately for each monkey. Filled bars, Monkey P; open bars, Monkey L. **C**) Learning rate ($\alpha$). **D**) Choice consistency ($\beta$). **E**) Probability of reward. **F**) Number of blocks performed. **G**) Engaged time per trial, estimated as the time between the appearance of the central target and the monkey touch of it . **H**) Response time, defined at the time between the stimuli onset and the detach (break) of the hand (gaze) from the central target (fixation point). Individual dots and lines represent individual sessions. Asterisks indicate significant differences between saccade and reach blocks (paired t-test; **** $p < 0.0001$; *** $p < 0.001$; ** $p < 0.01$; * $p < 0.05$; ns, not significant).

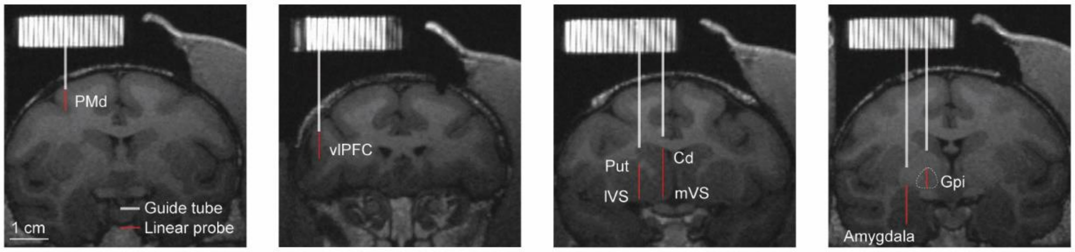


**Supplemental Figure 2**. **Recording sites.** Coronal MRI sections showing representative guide tube (white) and linear probe (red) trajectories for each targeted area: dorsal premotor cortex (PMd), ventrolateral prefrontal cortex (vlPFC), putamen (Put), inferior ventral striatum (IVS), caudate (Cd), medial ventral striatum (mVS), internal globus pallidus (GPi), and amygdala. Scale bar, 1 cm. Recording grid on top was filled with contrast agent during the MRI acquisition to allow planning of guide location and length, as well as the minimum and maximum depth of the tip of the linear probe.

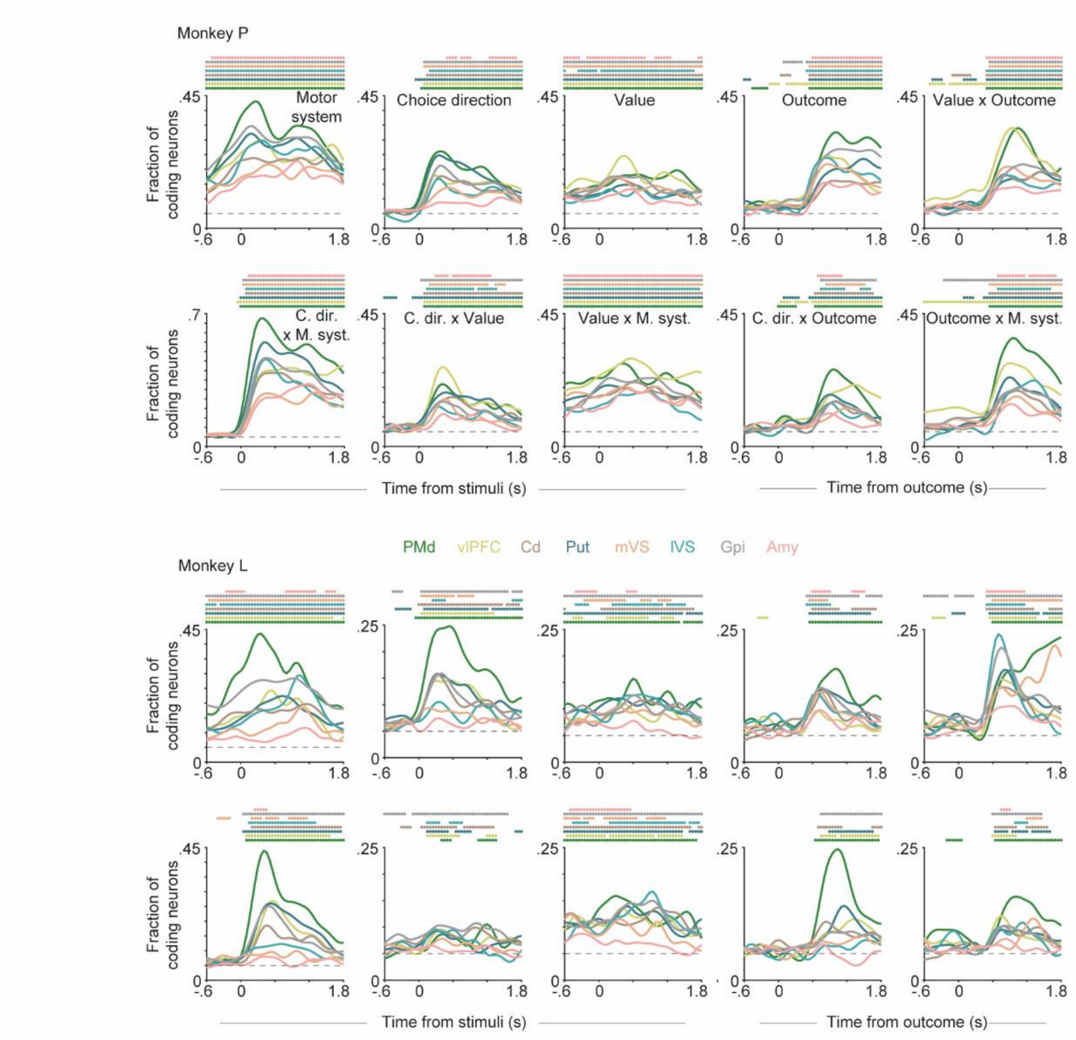


**Supplemental Figure 3**. **Full linear model results separated by monkey.** Fraction of neurons significantly encoding each main effect (motor system, choice direction, value, outcome) and interaction term over time, shown separately for Monkey P (top) and Monkey L (bottom). Left panels are aligned to stimuli presentation; right panels are aligned to outcome. Each colored line represents one area. Colored dots at top indicate time bins with fractions exceeding chance level (dashed line; binomial test, α = 0.05).

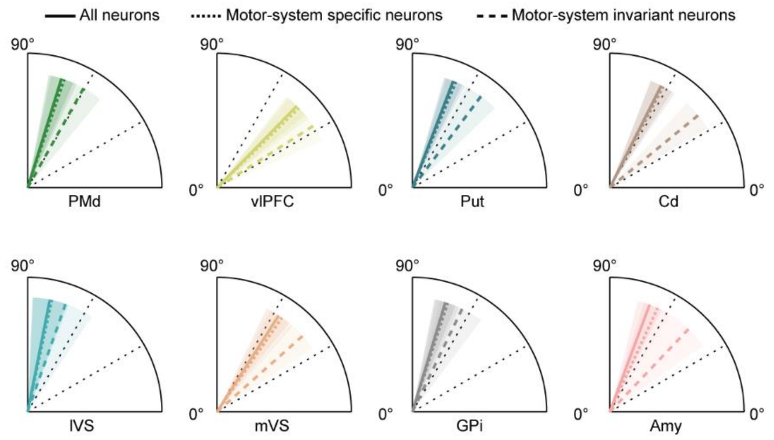


**Supplemental Figure 4**. **Pseudo-population angles by neuronal subpopulation.** Angle between saccade and reach value coding dimensions computed from pseudo-populations, shown separately for each area. Solid lines: all neurons. Dotted lines: embodied neurons only (significant value × motor system interaction). Dashed lines: motor-system-invariant neurons only (significant main effect of value). Shaded regions indicate ± SD across bootstrap iterations (n = 1000). Two-way ANOVA on the single value estimate per area and neuron coding subtype. Neuron coding subtype was significant: $F(2,23) = 53.737$, $p < 0.0001$. Mean angle ± SD across bootstraps: All neurons = 66.18° ± 10.58°, embodied neurons = 64.41° ± 10.70°, motor-system-invariant = 51.03° ± 13.22°. Post-hoc; neuron coding subtype pairwise comparisons (Bonferroni corrected): all neurons vs embodied neurons, $p = 0.862$; all neurons vs motor-system invariant, $p < 0.0001$; embodied neurons vs motor-system invariant, $p < 0.0001$.

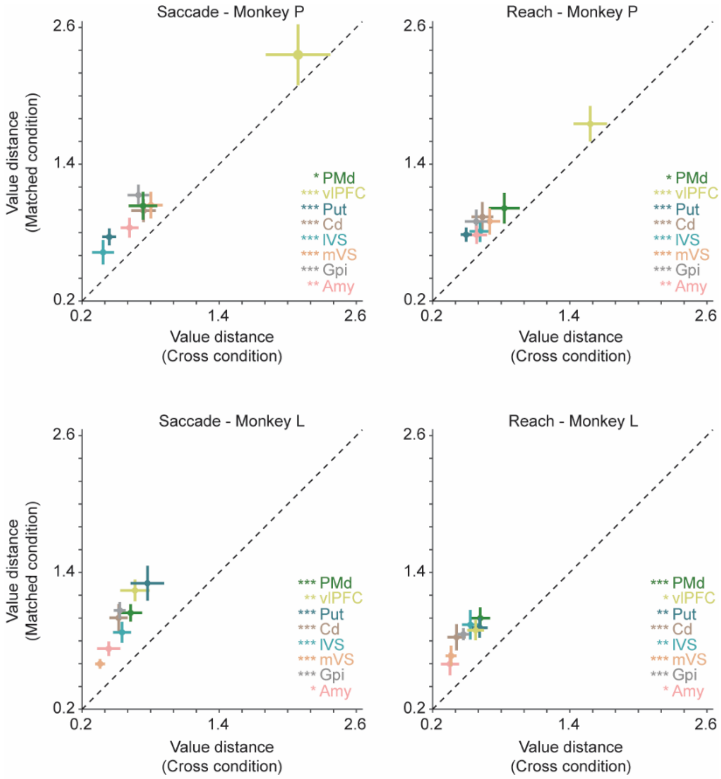


**Supplemental Figure 5. Cross-system value readout separated by monkey.** Within-system (y-axis) versus cross-system (x-axis) value distance during saccade (left) and reach (right) trials for Monkey P (top) and Monkey L (bottom). Each point represents the mean ± SEM across sessions for a given area. Dashed diagonal indicates equality. Points above the diagonal indicate stronger value discrimination under matched readout. Colored asterisks denote significant differences between matched and cross-system value distance (paired t-test, Bonferroni-corrected; * $p < 0.05$; ** $p < 0.01$; *** $p < 0.001$). The main effect of condition was significant in both monkeys for both saccade (Monkey P: $F(1,204) = 21.39$, $p < 0.0001$; Monkey L: $F(1,165) = 79.11$, $p < 0.0001$) and reach trials (Monkey P: $F(1,204) = 20.88$, $p < 0.0001$; Monkey L: $F(1,165) = 67.44$, $p < 0.0001$). The condition × area interaction was not significant in either monkey for saccade (Monkey P: $F(7,204) = 0.15$, $p = 0.99$; Monkey L: $F(7,165) = 0.99$, $p = 0.43$) or reach trials (Monkey P: $F(7,204) = 0.18$, $p = 0.99$; Monkey L: $F(7,165) = 0.28$, $p = 0.96$).

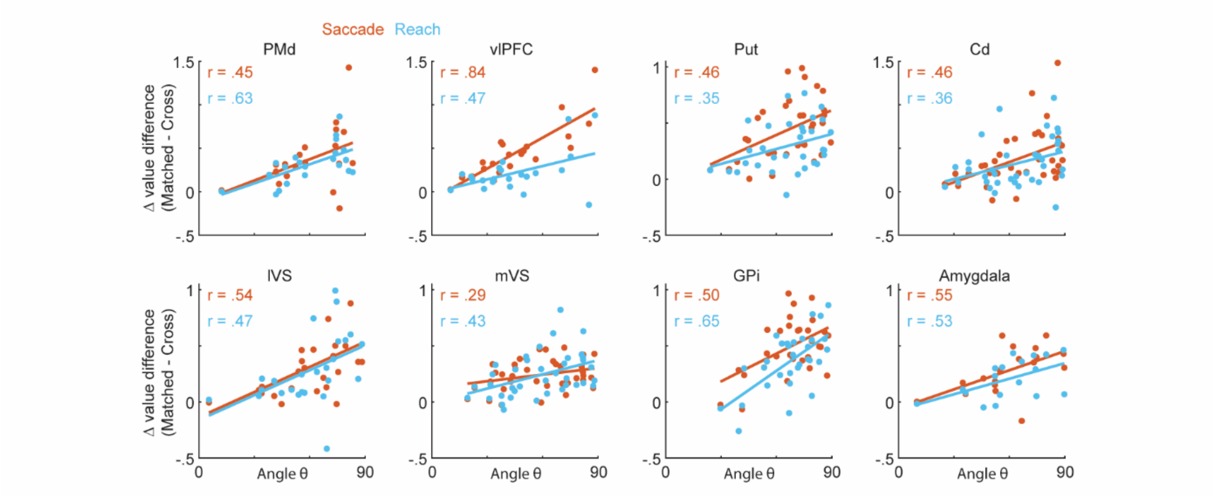


**Supplemental Figure 6. Correlation between geometric separation and value discriminability**. Pearson correlation between the angle (θ) separating saccade and reach value dimensions and the reduction in cross-system value discriminability (Δ value distance: matched − cross) across sessions, shown separately for each area. Each dot represents one session. Orange: saccade trials. Blue: reach trials. Correlation coefficients (r) are reported for each area and motor system.

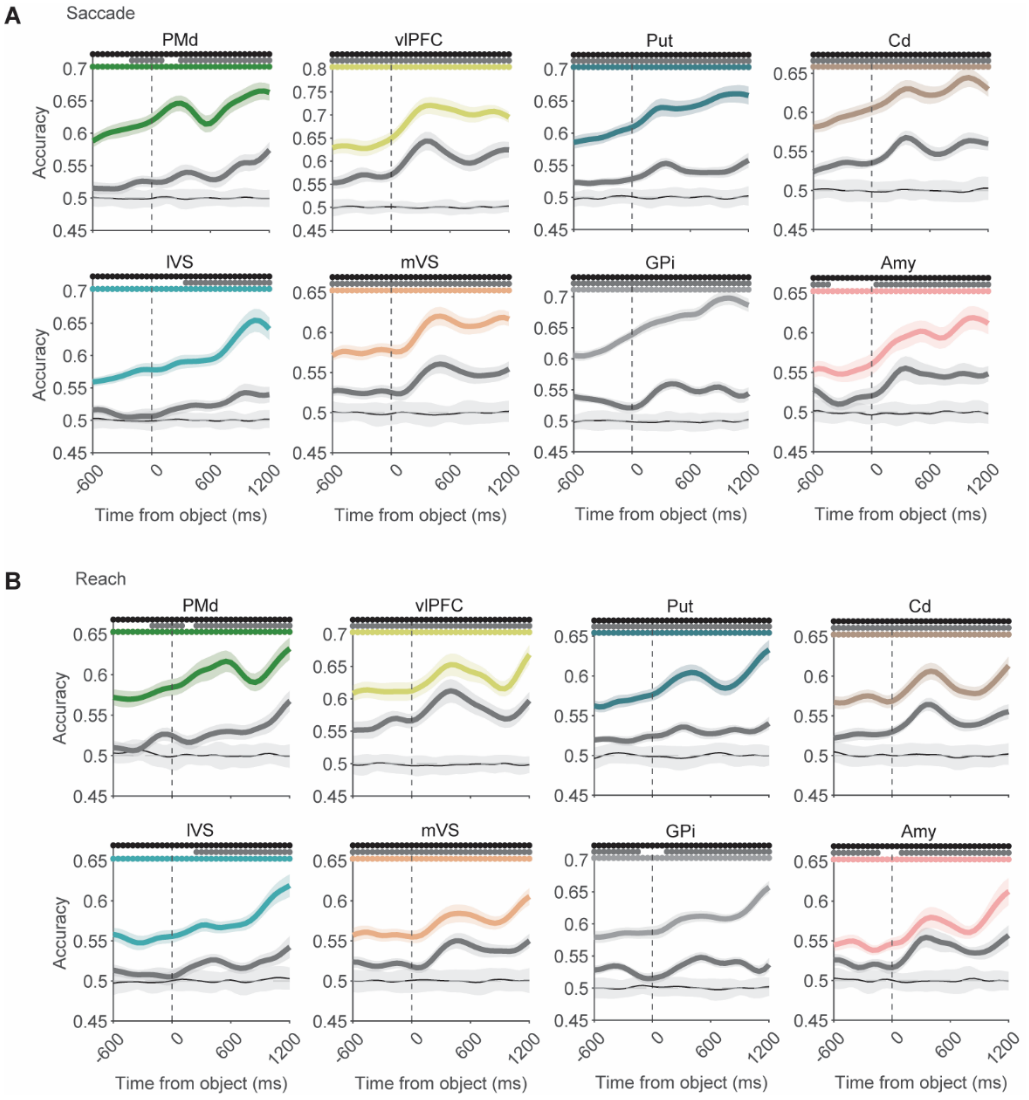


**Supplemental Figure 7. Time-resolved within-system and cross-system decoding of value. A)** Saccade trials. **B)** Reach trials. For each area, colored lines show within-system decoding accuracy and dark gray lines show cross-system decoding accuracy over time (mean ± SEM across sessions). Light gray lines and shading indicate chance level estimated from label-shuffled null distributions. Dashed vertical lines indicate stimuli presentation. Colored and gray dots at top indicate time bins with within-system and cross-system accuracy significantly exceeding chance, respectively. Black dots indicate time bins with significance difference between within-system and cross-system accuracy. Paired t- test ($p < 0.05$).

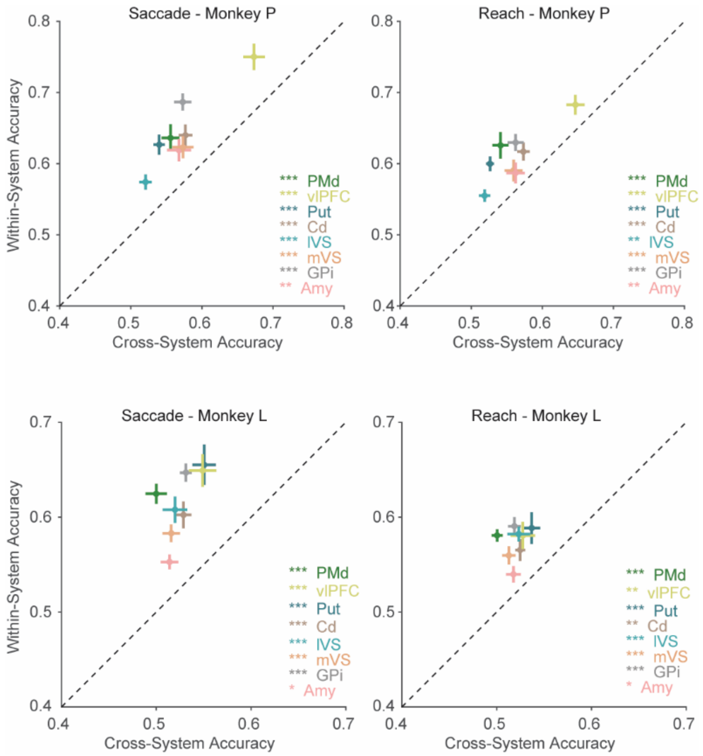


**Supplemental Figure 8. Cross-system decoding separated by monkey.** Within-system (y-axis) versus cross-system (x-axis) decoding accuracy for saccade (left) and reach (right) trials, shown separately for Monkey P (top) and Monkey L (bottom). Each point represents the mean ± SEM decoding accuracy for a given area, averaged during the 600 ms following stimuli presentation. Dashed diagonal indicates equal accuracy. Colored asterisks denote significant differences between within-system and cross-system accuracy (paired t-test, Bonferroni-corrected; * $p < 0.05$; ** $p < 0.01$; *** $p < 0.001$). The main effect of condition was significant in both monkeys for both saccade (Monkey P: $F(1,204) = 119.32$, $p < 0.0001$; Monkey L: $F(1,165) = 247.26$, $p < 0.0001$) and reach trials (Monkey P: $F(1,204) = 83.13$, $p < 0.0001$; Monkey L: $F(1,165) = 67.44$, $p < 0.0001$).

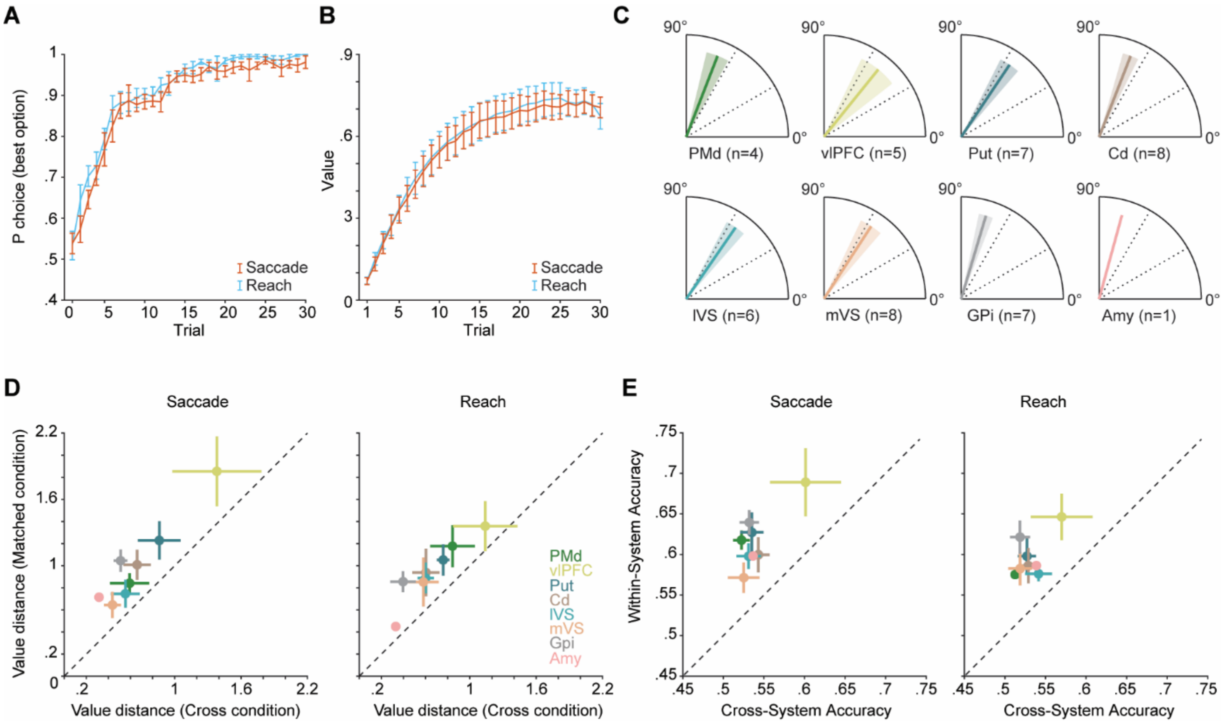


**Supplemental Figure 9. Embodied organization under matched learning performance and value estimate.** A) Learning curves and B) stimulus-value estimates for saccade (orange) and reach (blue) blocks in the subset of sessions with similar behavior between motor systems (n = 8 sessions; paired t-test, $p > 0.05$). C) Angle between saccade and reach value coding dimensions for each area in the matched-behavior subset. Angles from behaviorally similar sessions remained largely orthogonal (79.43° ± 4.77°). D) Within-system versus cross-system value distance. In the full dataset, the condition × behavioral similarity interaction was not significant for either saccade ($F(1,429) = 0.16$, $p = 0.69$) or reach trials ($F(1,429) = 0.08$, $p = 0.78$), while the main effect of condition remained significant for both (saccade: $F(1,429) = 36.33$, $p < 0.0001$; reach: $F(1,429) = 37.91$, $p < 0.0001$). In the behaviorally similar subset, the main effect of condition remained significant (saccade: $F(1,69) = 5.82$, $p = 0.019$; reach: $F(1,69) = 6.38$, $p = 0.014$), with no condition × area interaction (saccade: $F(7,69) = 0.16$, $p = 0.99$; reach: $F(7,69) = 0.11$, $p = 0.99$). E) Within-system versus cross-system decoding accuracy. In the full dataset, the condition × behavioral similarity interaction was not significant for either saccade ($F(1,429) = 0.24$, $p = 0.62$) or reach trials ($F(1,429) = 2.90$, $p = 0.09$), while the main effect of condition remained significant for both (saccade: $F(1,429) = 144.14$, $p < 0.0001$; reach: $F(1,429) = 108.02$, $p < 0.0001$). In the behaviorally similar subset, the main effect of condition remained significant (saccade: $F(1,69) = 35.55$, $p < 0.0001$; reach: $F(1,69) = 34.43$, $p < 0.0001$), with no condition × area interaction (saccade: $F(7,69) = 0.63$, $p = 0.73$; reach: $F(7,69) = 0.68$, $p = 0.69$). Conventions as in main Figures 1, 3-5.

**Supplemental Table 1.** Number of neurons recorded in each session and area separately for monkey P (MP) and monkey L (ML)

| Session-Area | PMd | vlPFC | Put | Cd | lVS | mVS | GPi | Amy |
|---|---|---|---|---|---|---|---|---|
| 1 (MP) | | | 33 | 10 | 23 | 4 | 21 | |
| 2 (MP) | | | 43 | 15 | 16 | 5 | 10 | |
| 3 (MP) | | | 35 | 12 | 4 | 12 | | |
| 4 (MP) | 8 | | 42 | 39 | 20 | 24 | 24 | |
| 5 (MP) | 17 | | 11 | 24 | 19 | 16 | 25 | |
| 6 (MP) | 24 | | 26 | 34 | 16 | 23 | 28 | |
| 7 (MP) | 2 | | 40 | 15 | 18 | 22 | 32 | |
| 8 (MP) | 31 | | 20 | 16 | 11 | 14 | 38 | |
| 9 (MP) | | 38 | 42 | 11 | | 11 | 32 | |
| 10 (MP) | | 22 | 37 | 14 | | 8 | 33 | |
| 11 (MP) | | 34 | 19 | 12 | | 12 | 33 | |
| 12 (MP) | | 23 | 13 | 10 | | 16 | 31 | |
| 13 (MP) | | 12 | 17 | 11 | | 8 | 31 | |
| 14 (MP) | | 15 | 14 | 3 | | 2 | 29 | |
| 15 (MP) | 13 | 20 | 8 | 10 | 4 | 17 | | |
| 16 (MP) | 28 | 4 | 6 | 16 | 10 | 11 | | |
| 17 (MP) | 26 | 25 | 8 | 34 | 10 | 15 | | |
| 18 (MP) | 30 | 22 | 8 | 27 | | 24 | | |
| 19 (MP) | 34 | 26 | 18 | 29 | 2 | 15 | | |
| 20 (MP) | 27 | 20 | 16 | 27 | 3 | 17 | | |
| 21 (MP) | 30 | 25 | 3 | | 5 | 4 | | |
| 22 (MP) | | | | | | | | 8 |
| 23 (MP) | | | | | | | | 16 |
| 24 (MP) | | | | | | | | 3 |
| 25 (MP) | | | | | | | | 32 |
| 26 (MP) | | | | | | | | 39 |
| 27 (MP) | | | | | | | | 22 |
| 28 (MP) | | | | | | | | 30 |
| 29 (MP) | | | | | | | | 47 |
| 30 (MP) | | | | | | | | 31 |
| 31 (MP) | | | | | | | | 20 |
| 32 (MP) | | | | | | | | 8 |
| 33 (ML) | | | | | | | 31 | 20 |
| 34 (ML) | | | 34 | 13 | 19 | 7 | 31 | |
| 35 (ML) | | | 35 | 27 | 15 | 18 | 27 | |
| 36 (ML) | | | | 9 | | 9 | 20 | 44 |
| 37 (ML) | | | | 33 | | 20 | 29 | 64 |
| 38 (ML) | | | | 28 | | 21 | 29 | 42 |
| 39 (ML) | | | | 39 | | 18 | 37 | 54 |
| 40 (ML) | | | | 27 | | 19 | 26 | 38 |
| 41 (ML) | | | | 32 | | 13 | 21 | 30 |
| 42 (ML) | 17 | 23 | 45 | 29 | 18 | 12 | 17 | |
| 43 (ML) | 21 | 32 | 34 | 35 | 24 | 17 | 31 | |
| 44 (ML) | 30 | 33 | 45 | 10 | 28 | 12 | 31 | |
| 45 (ML) | 19 | 31 | 35 | 11 | 10 | 17 | 29 | |
| 46 (ML) | 25 | 31 | 39 | 23 | 16 | 13 | 31 | |
| 47 (ML) | 20 | 29 | 7 | 13 | 4 | 12 | 13 | |
| 48 (ML) | 30 | 31 | 18 | 16 | 6 | 17 | 24 | |
| 49 (ML) | 30 | 30 | 20 | 6 | 15 | 3 | 26 | |
| 50 (ML) | 31 | 30 | 26 | 2 | 18 | 2 | 27 | |
| total MP | 270 | 286 | 459 | 369 | 161 | 280 | 367 | 256 |
| total ML | 223 | 270 | 338 | 353 | 173 | 230 | 480 | 292 |
| total (MP+ML) | 493 | 556 | 797 | 722 | 334 | 510 | 847 | 584 |
